# Low expression of ANT1 confers oncogenic properties to rhabdomyosarcoma tumor cells via modulating metabolism and death pathways

**DOI:** 10.1101/2020.03.03.973719

**Authors:** J Vial, P Huchedé, S Fagault, F Basset, M Rossi, J Geoffray, J Bisaccia, M Creveaux, D Neves, F Fauvelle, P Castets, M Carré, K Weber, M Castets

**Affiliations:** Cell death and Childhood Cancers Laboratory – Equipe labellisée LabEx DEV2CAN, Centre de Recherche en Cancérologie de Lyon, INSERM U1052-CNRS UMR5286, Université de Lyon, Centre Léon Bérard, 69008, Lyon, France; Netris Pharma, Lyon, France; Aix-Marseille Université, Inserm UMR_S 911, Centre de Recherche en Oncologie biologique et Oncopharmacologie, Faculté de pharmacie, Marseille, France; University Grenoble Alpes, INSERM, US17, MRI facility IRMaGe, 38000 Grenoble, France; Department of Cell Physiology and Metabolism, Université de Genève, CMU, CH-1211 Genève, Switzerland

## Abstract

Rhabdomyosarcoma (RMS) is the most frequent form of pediatric soft-tissue sarcoma. It is divided into 2 main subtypes: ERMS (embryonal) and ARMS (alveolar). Current treatments are based on chemotherapy, surgery and radiotherapy. 5-year survival rate remains of 70% since 2000, despite several clinical trials.

RMS cells are thought to derive from muscle lineage precursors. During development, myogenesis is characterized by primary expansion of myoblasts, elimination of those in excess by cell death and the differentiation of the remaining ones into myotubes and myofibers. The idea that these processes could be hijacked by tumor cells to sustain their oncogenic transformation has emerged, while RMS is being considered as the Mister Hyde’s side of myogenesis. Thus, focusing on myogenic developmental programs could help understanding RMS molecular aetiology.

Following this idea, we decided to concentrate on ANT1, which is involved in myogenesis and is the underlying cause of genetic disorders associated with muscle degeneration. ANT1 is a mitochondrial protein, which has a functional duality, as it is involved both in metabolism via regulation of ATP/ADP release from mitochondria, but also in apoptosis as part as the mitochondria Permeability Transition Pore (mPTP). By bioinformatic analysis of transcriptomic datasets, we observed that ANT1 is expressed at low levels in RMS. Using CRISPR-Cas9 technology, we showed that decreased ANT1 expression confers selective advantages to RMS cells in terms of proliferation and resistance to stress-induced death. These effects result notably from a metabolic switch. Restoration of ANT1 expression using a Tet-On system is sufficient to prime tumor cells to death and to increase their sensitivity to chemotherapies. Thus, modulation of ANT1 activity could appear as an appealing therapeutic approach in RMS management.

## INTRODUCTION

Childhood cancers may be considered as aberrations of pre- and postnatal ontogeny, resulting from dysregulated developmental processes (1, 2). Development is characterized by rapid proliferative expansion of cells, their migration along proper routes towards appropriate organs, tissue refinement by elimination of excessive cells, and the terminal differentiation of survivors into proper cell types.

In particular, during development, cells in excess are secondarily pruned through switches in developmental programs governing survival: the regulation of cell suicide pathways is then crucial. It is well known for example that during early embryogenesis, a pool of myoblasts is generated in somites during skeletal muscle development (3): a fraction of these cells will undergo differentiation, another subset will exit the cell cycle to constitute a niche of resting stem cells, while the remaining will be eliminated by apoptosis (3). In some rare instances, some supernumerary cells can resist these death signals as a potential first pathological step towards cancer (1). Indeed, resistance to cell death plays a key role in the first pathological steps towards cancer as well as in constitutive or acquired resistance to treatments (4).

Rhabdomyosarcoma (RMS) is the most frequent form of paediatric soft tissue sarcoma, accounting for 5% of solid paediatric tumors, with a 5-years survival that caps at 60-70% (5, 6). Molecular basis of RMS remains unclear, notably in the sizeable fraction of translocation negative tumors. Regarding cell death resistance, overall survival has been correlated with the expression of Bcl family members in patients and resistance to apoptosis linked to failure of conventional therapies in RMS cell lines. However, robust characterization of cell death pathways altered in this cancer is still lacking (5, 7). RMS are thought to arise from malignant transformation of muscle precursors (6). Embryonal RMS (ERMS), so-called because they resemble embryonic muscle, present various degree of myogenic differentiation, ranging from small round cells to larger oblong ones, sometimes having a strap-like appearance for the most differentiated ones, occasionally with cross striations and multinucleation (6). Besides these morphological observations supporting myoblast-like nature of tumor cells, RMS expression signatures are also characterized by widespread expression of embryonic muscle specific markers such as Myogenin, Desmin or MyoD. Moreover, the notion that molecular pathogenesis of RMS could share similarities with processes involved in myogenesis or that have gone awry in muscular dystrophies has begun to emerge (8–10).

Besides gold standard effectors of death pathways, we then made the hypothesis that genes involved in specific cell survival/death imbalance during myogenesis or in muscular dystrophies etiology could also play a role in RMS tumorigenesis.

Based on exhaustive bioinformatics-driven bibliographic data mining, we decided to focus on the role that ANT1 (Adenine Nucleotide Translocator 1) could play in these cancers. ANT1 is the heart and muscle-specific isoform of the ANT mitochondrial inner membrane proteins family, and has been associated to mitochondrial myopathy and cardiomyopathy (11–13). It has also been involved in myoblast differentiation (14). ANT1 regulates ATP/ADP exchange across the mitochondrial inner membrane. Indeed, it exports ATP produced by OXPHOS metabolism in the cytoplasm to power cellular reactions and imports simultaneously ADP to restore intra-mitochondrial stock. ANT1 also interacts with VDAC proteins in the so-called mPTP, whose opening triggers mitochondrial membrane permeabilization, release of cytochrome C and subsequent activation of intrinsic apoptotic pathway (15–17). ANT1 is then considered to be at the crossroad of several cell death and metabolic signalling paths. Here, we show that ANT1 is expressed at low levels in RMS tumors. Using inducible CRISPR-Cas9 strategy, we observed that down-regulation of ANT1 expression level expression increases both cell proliferation and resistance to stress-induced cell death, the last being counteracted by restoration of ANT1 expression. Thus, ANT1 behaves as a new tumor suppressor in RMS.

## RESULTS

### Low level of ANT1 expression in RMS favors tumor cell proliferation

Alterations of ANT1 expression level have been reported in several disorders and notably associated with muscular defects (18–21). However, its expression profile in cancers has almost not been studied so far, especially in RMS. To analyse the expression level of ANT1 in RMS, we first performed a bioinformatic analysis of the publicly available GSE28511 dataset, dedicated to the comparison of normal skeletal muscle tissue (n=6) versus RMS (n=8 ERMS and n=10 ARMS). We observed that the expression of ANT1 encoding gene, *SLC25A4*, is significantly lower in RMS than in non-tumoral skeletal muscle: its expression is decreased by 24 folds, as compared to only 3 folds for the electron transport chain protein COX7C or the mitochondrial hexokinase HK1 for example (Figure 1A). Reciprocally, the level of expression of ANT2 encoding gene, *SLC25A5*, is increased by 3 folds in tumors, as previously reported in other cancers (22, 23). We confirmed this result by Q-RT-PCR, by showing that ANT1 is more expressed in adult, fetal and dystrophic muscles than in 67 pediatric RMS samples (Figure 1B). Using E-TABM-1202, we observed that high ANT1 expression level may be positively associated to a better outcome in fusion-negative RMS, also this observation needs to be confirmed on a higher number of patients (Figure S1A). We hypothesized that low ANT1 expression level may confer a selective advantage to tumor cells. In patients, we observed using R2 cancer software that ANT1 expression tends to be negatively correlated to CDC7, which has been described as an inducer of smooth muscle cell proliferation (24), and to Ki67 suggesting that ANT1 may impact cell proliferation (Figure S1B-C). To investigate if ANT1 impacts cell proliferation, we screened its expression in different RMS cell lines by Q-RT-PCR (Figure 1C). We chose the ERMS CCL-136 cell line, which has the highest level of ANT1 expression, to study impact of ANT1 knock-down. We set up a stable doxycycline-inducible CRISPR-Cas9 system that leads to a reduction in ANT1 expression of 80% (Figure 1D), without any impact on ANT2 expression (Figure S1D). These cells will be further referred as CCL-136^Low^. Decrease in ANT1 expression in these cells is associated to an increase in cell density by 2.8, 2.1 and 1.85, 24h, 48h and 72h post-silencing as measured by WST1 assay (Figure 1E), and to a 42.5% increase of viable cells concentration in normal growth conditions (Figure S1E). Similar result was observed after silencing of ANT1 by siRNA in immortalized myoblasts (Figure S1F-G). Thus, RMS are associated with low level of ANT1 expression, which could sustain tumor cells proliferation.

**Figure 1:**
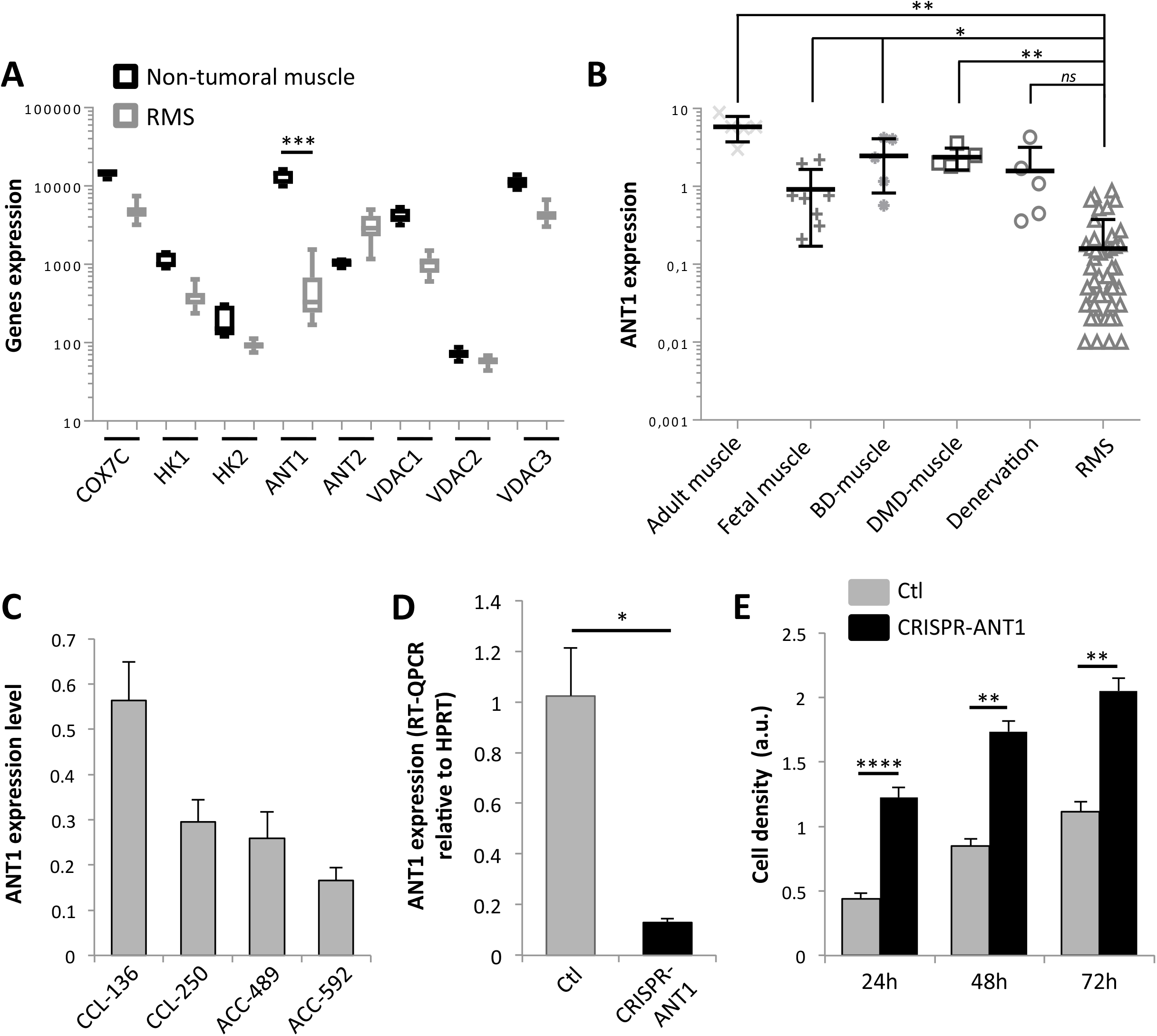
Low level of ANT1 expression in RMS favors tumor cell proliferation. **a.** Bioinformatic analysis of the expression of ANT1 gene (*SLC25A4*) and of some of its partners in a cohort of non-tumoral muscle (n=6), ERMS (n=8) and ARMS (n=10) from the GSE28511 transcriptomic dataset. ***: p<0.001, two-sided independent samples T-test. **b.** Quantification of ANT1 gene (*SLC25A4*) expression by Q-RT-PCR, relatively to *HPRT* housekeeping gene in RMS biopsies (n=67), normal adult muscle (n=5), fetal muscle (n=9), and in biopsies from patients with muscle weakness (BD: Becker Dystrophy, DMD: Duchenne Muscular Dystrophy, n=5 each). *: p<0.05, **: p<0.01, ***: p<0.001, *ns*: not significant; two-sided independent samples T-test. **c.** Quantification of ANT1 expression by Q-RT-PCR, relatively to *HPRT* housekeeping gene in ERMS cell lines (CCL-136 and CCL-250), and ARMS cell lines (ACC-489 and ACC-592). Results are presented as means+/−std; n=3. **d.** Efficiency of ANT1 silencing by CRISPR-Cas9 in CCL-136 cells, 48 hours after doxycycline treatment. Quantification of ANT1 (*SLC25A4*) expression by Q-RT-PCR, relatively to *HPRT* housekeeping. Results are presented as means+/−std; n=3. *: p<0.05; independent samples T-test. **e.** Increase in cell density as measured by WST1 assay in ANT1^Low^ CCL-136 cells, at different time points after silencing of ANT1 expression by doxycycline treatment. Results are presented as means+/−std; n=3. **: p<0.01, ****: p<0.0001; two-sided independent samples T-test.

### Loss of ANT1 confers selective advantage to tumor cells by maintaining them in a proliferative state

ANT1 is at the crossroad of metabolic, death but also mitogen-activated signals (25). Thus, increase in cell density observed in CCL-136^Low^ could have multiple origins. Since ANT1 is known to play a role in metabolic regulation through control of ATP/ADP exchange, we hypothesized that ANT1 decrease be associated with some metabolic changes. Metabolomic analysis revealed significant decrease in amino-acids content in CCL-136^Low^, notably impacting glycine (Figure 2A). Amino acids especially those linking to tricarboxylic acid cycle are an alternative energy source used during cancer cell proliferation (26). Consistently, such decrease in glycine content has previously been associated with increased proliferation rate in cancer cells (27). It has been shown that differentiation of myoblasts is associated with decrease in phosphocholine content (28). The levels of *scyllo*-Inositol, Choline and Glycerophosphocholine are on the contrary significantly increased in CCL-136^Low^ (Figure 2B). Such increases are reminiscent of activation of choline metabolism, which is a hallmark of cancer cells (29). Export of ATP is coupled to the functioning of the multi-proteic complexes of the electron transport chain, driving OXPHOS metabolism. Using Seahorse system, we showed that decrease of ANT1 expression is also associated with a significant increase in basal oxygen consumption rate (OCR, Figure 2C), as a marker of OXPHOS metabolism. Such increase in OXPHOS metabolism has been described to occur upon myogenic precursors activation during regeneration (30). CCL-136^Low^ were slightly less glycolytic with a basal extracellular acidification (ECAR) decrease of 31% as compared to controls (Figure 2D). The spare respiratory capacity (SRC) measures the extra mitochondrial capacity available in a cell to produce energy under conditions of increased demand and is calculated by the difference between the maximal and basal OCR (Figure 2C and Material & Methods). SRC remains unchanged when ANT1 level is lowered, indicating that those cells are still able to adapt to metabolic stress.

**Figure 2 :**
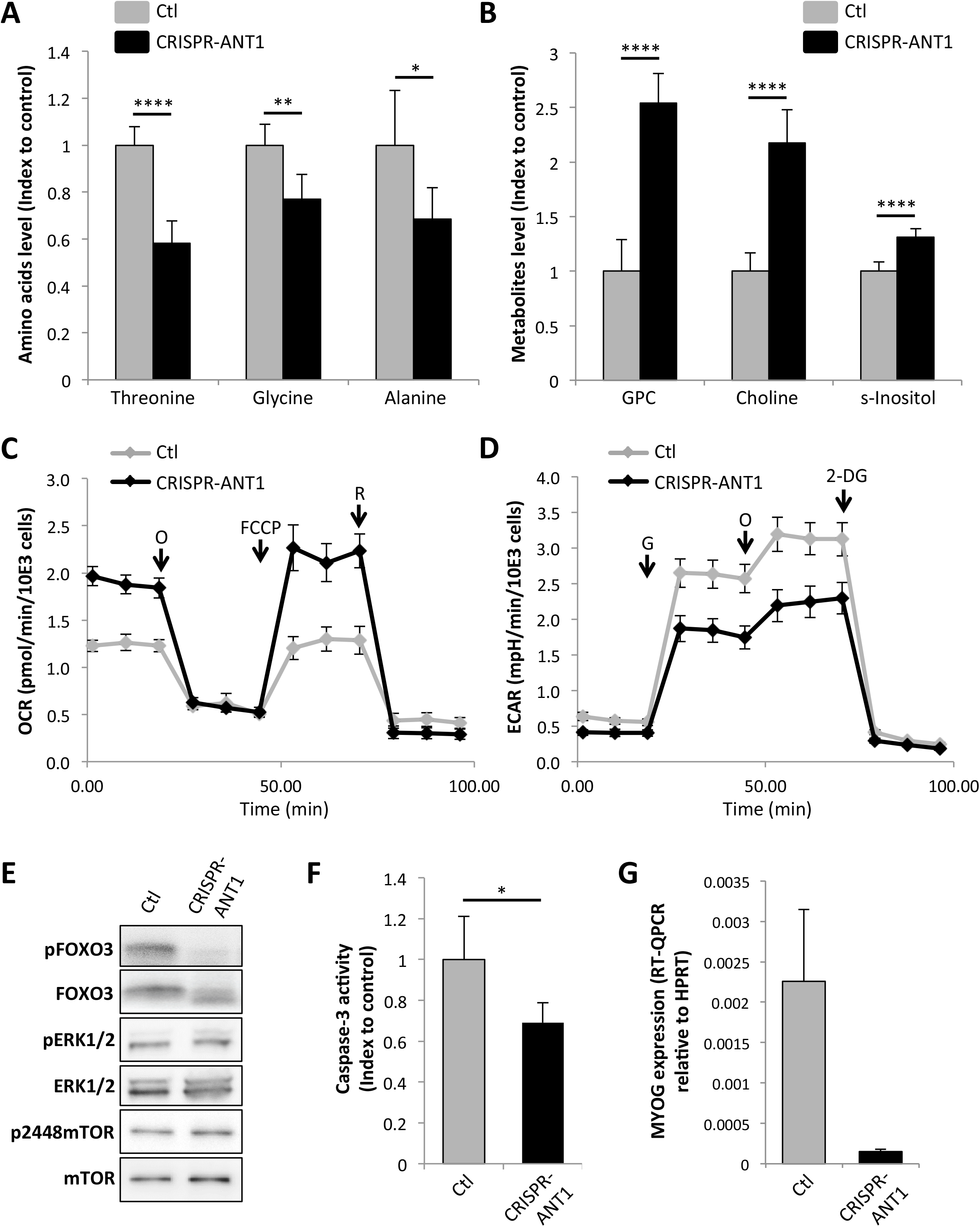
Loss of ANT1 confers selective advantage to tumor cells by maintaining them in a proliferative state. **a.** Changes in amino acids abundance in ANT1^Low^ CCL-136 cells as compared to control ones. Results are presented as means+/−std of the ratio between metabolites abundance in ANT1^Low^ CCL-136 cells as compared to control ones; n=5. *: p<0.05, **: p<0.01, ****: p<0.0001; two-sided independent samples T-test. **b.** Changes in choline derivatives and inositol metabolites abundance in ANT1^Low^ CCL-136 cells as compared to control ones. GPC: Glycerophosphocholine, s-Inositol: *scyllo*-Inositol. Results are presented as means+/−std of the ratio between metabolites abundance in ANT1^Low^ CCL-136 cells as compared to control ones; n=5. p<0.0001; two-sided independent samples T-test. **c.** Oxygen Consumption Rate (OCR) measured by Seahorse in ANT1^Low^ CCL-136 cells as compared to control ones. O: Oligomycin, FCCP: Carbonyl cyanide-4-(trifluoromethoxy)-phenylhydrazone, R: Rotenone. Results are presented as means+/−std; n=3. **d.** Extracellular Acidification Rate (ECAR) in ANT1^Low^ CCL-136 cells as compared to control ones, as a hallmark of glycolysis. G: Glucose, O: Oligomycine, 2-DG: 2-deoxy-glucose. Results are presented as means+/−std; n=3. **e.** Decrease of FOXO3 and of activated pFOXO3 levels in ANT1^Low^ CCL-136 cells. ANT1 loss does seem to have any impact on activation of ERK1/2 and mTOR pathways, evaluated by phosphorylation level on Western Blots. Tubulin is used as a loading control. **f.** Increase in basal level of caspase-3 activity in ANT1^Low^ CCL-136 cells. Results are presented as means+/−std; n=3. *: p<0.05; two-sided independent samples T-test. **g.** Decrease in the expression of commitment marker Myogenin in ANT1^Low^ CCL-136 cells as compared to control ones. Results are presented as means+/−std of 3 independent Q-RT-PCR.

Beside this metabolic switch and since ANT1 was notably reported to modulate ERK1/2 activation (25), we assessed the status of key pathways involved in regulation of myoblasts proliferation. Whereas activation levels of ERK1/2 and mTOR do not appear to be modified (Figure 2E), we observed that FOXO3 level, which has been demonstrated to impair muscle progenitors proliferation (31, 32), is significantly decreased in CCL-136^Low^ (Figure 2E). During myogenesis, it was shown that the caspases, and notably caspase-3, activation is required at a ‘sub-apoptotic’ level to engage myoblasts in differentiation (33–35). Caspases are cysteine proteases executioners of apoptosis. Since ANT1 was shown to regulate apoptosis (36, 37), we wondered whether modulation of its expression could also have an impact on this death-associated signalling cascade. Indeed, we observed that basal level of caspase-3 activity is significantly decreased in CCL-136^Low^ (Figure 2F). Consistently, we observed that expression of the commitment marker Myogenin is significantly reduced in these cells as compared to control ones (Figure 2G). ANT1 expression level tends to be also negatively correlated with specific skeletal muscle-genes markers such as Troponin 3 and MYBPC1 in patients, suggesting an association between ANT1 expression level and tumor cell differentiation status (R2 cancer analysis, GSE16476, Figure S2A-B).

Altogether, these observations suggest that low level of ANT1 expression is sufficient to maintain RMS tumor cells in an immature proliferative state by acting on different cellular parameters.

### Low levels of ANT1 favors resistance of tumor cells to death induced by stress and chemotherapies

ANT1 was shown to exert a pro-death function notably in conditions of oxidative stress response (38), as part of the mPTP. We then decided to study impact of modulation of its expression level in response to H_2_O_2_ treatment using our CRISPR-Cas9 system. Silencing of ANT1 is associated with an increase in the percentage of viable CCL-136^Low^, as compared to controls (Figure 3A). Consistently, this effect is accompanied by a loss of mitochondrial membrane potential (Δψ_m_), indicative for the induction of intrinsic apoptotic pathway (Figure 3B). H_2_O_2_ triggers death via generation of reactive oxygen species (ROS). Increase in cell viability in CCL-136^Low^ is accompanied by a decrease in ROS production, as measured by follow up of CellROX®, suggesting that detoxification of ROS is more efficient in these ANT1 silenced cells (Figure 3C).

**Figure 3:**
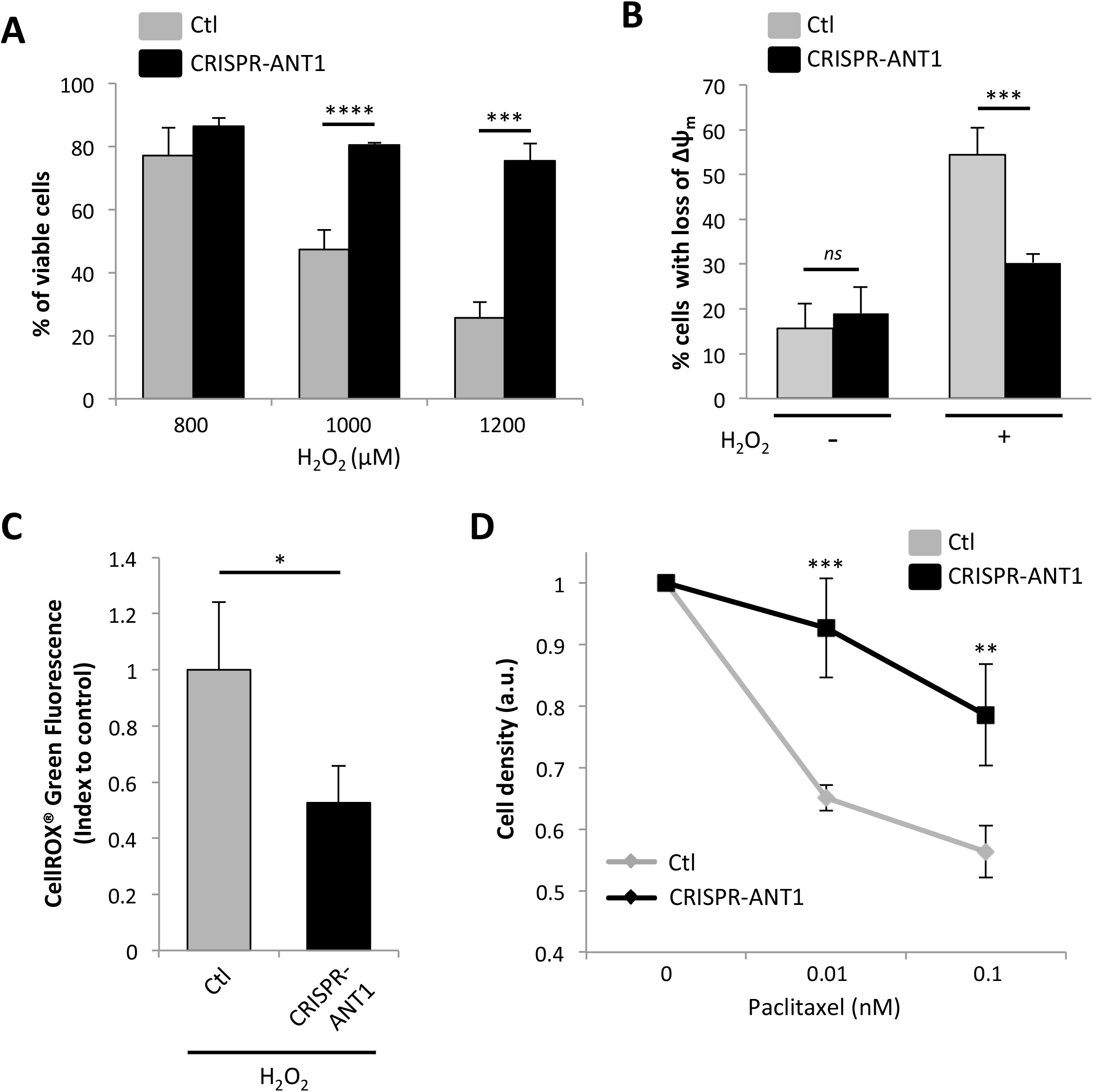
Low levels of ANT1 favors resistance of tumor cells to death induced by stress and chemotherapies. **a.** Increase in percentage of viable cells in ANT1^Low^ CCL-136 cells treated upon H_2_O_2_, treatment. Results are presented as means+/−std; n=3. ***: p<0.001, ****: p<0.0001; two-sided independent samples T-test. **b.** Decrease of the cell percentage with loss of mitochondrial membrane depolarization (Δψ_m_) in ANT1^Low^ CCL-136 cells upon H_2_O_2_ stress, compared to control cells. Results are presented as means+/−std; n=3. ***: p<0.001, *ns*: not significant; two-sided independent samples T-test. **c.** ROS production is lower in ANT1^Low^ CCL-136 cells upon H_2_O_2_ stress. ROS production was evaluated by measure of CellROX® fluorescence. Results are presented as means+/−std; n=3. *: p<0.05; two-sided independent samples T-test. **d.** Decreased sensitivity of ANT1^Low^ CCL-136 cells to Paclitaxel after 24 hours of treatment, compared to control cells (WST1 assay). Results are presented as means+/−std; n=3. **: p<0.01, ***: p<0.001; two-sided independent samples T-test.

We then assessed impact of ANT1 low expression level on modulation of sensitivity to chemotherapies. As shown on Figure 3D, decrease of ANT1 expression is associated with a reduced sensitivity to chemotherapies treatments, as exemplified with Paclitaxel (Figure 3D). Same results were obtained with vincristine (not shown).

Thus, beside promotion of cell proliferation, low level of ANT1 is also sufficient to increase the resistance of RMS tumor cells to death, thereby conferring a second selective advantage to these cells.

### Restoration of ANT1 is sufficient to sensitize RMS cells to death induced by stress and chemotherapies

Loss of ANT1 seems to confer selective advantages to RMS tumor cells. We then wondered whether restoration of its expression may have tumor suppressor effects. We then designed Tet-On inducible stable CCL-136 and ACC-489 cell lines to define the impact of restoration of ANT1 expression on stress-induced tumor cell death (Figures S3A-F). These cells will be further designed as CCL-136^High^ and ACC-489^High^. Since ACC-489 presents the lowest level of ANT1 expression, we chose to study the impact of ANT1 expression restoration preferentially in these cells. It was shown that ANT1 overexpression drives apoptotic cell death, probably resulting from aberrant permeabilization of mitochondrial membrane (36). We chose to use a doxycycline dose that was sufficient to restore ANT1 expression without impact on tumor cell death promotion in normal growth conditions (Figure 4A). In this steady-state condition, we however observed an increase in caspase-3 activity in ACC-489^High^ cells, independently of any death stimuli (Figure 4B). Consistently with the observation that cell viability was not affected by restoration of ANT1 expression, we do not however detect any impact of this caspase-3 activation on PARP cleavage, as a downstream effector of intrinsic apoptotic pathway, in these basal conditions (Figure 4C). We then wondered whether those cells may be more sensitive to stress. Restoration of ANT1 expression is indeed sufficient to drive increased sensitivity to H_2_O_2_ treatment both in ACC-489 ^High^ and CCL-136^High^ cell lines (Figure 4D and Figure S3G). Increase in cell death in ACC-489 ^High^ upon H_2_O_2_ treatment is accompanied by an increase in the percentage of cells with mitochondrial membrane depolarisation (Figure 4E). Beside oxidative stress, we also observed that viability of ACC-489^High^ is reduced upon chemotherapies treatments, as exemplified with Paclitaxel (Figure 4F and Figure S3H) and Vincristine (Figure 4G). To go one step further, we set up RMS 3D-spheroid models using both ACC-489 control and ACC-489^High^ RMS cells. We observed that re-expression of ANT1 in this 3D-system is also sufficient to decrease resistance to chemotherapies (Figure 4I, left panel), likely via apoptosis induction since it is associated with an increase in caspase-3 activity (Figure 4J, right panel).

**Figure 4:**
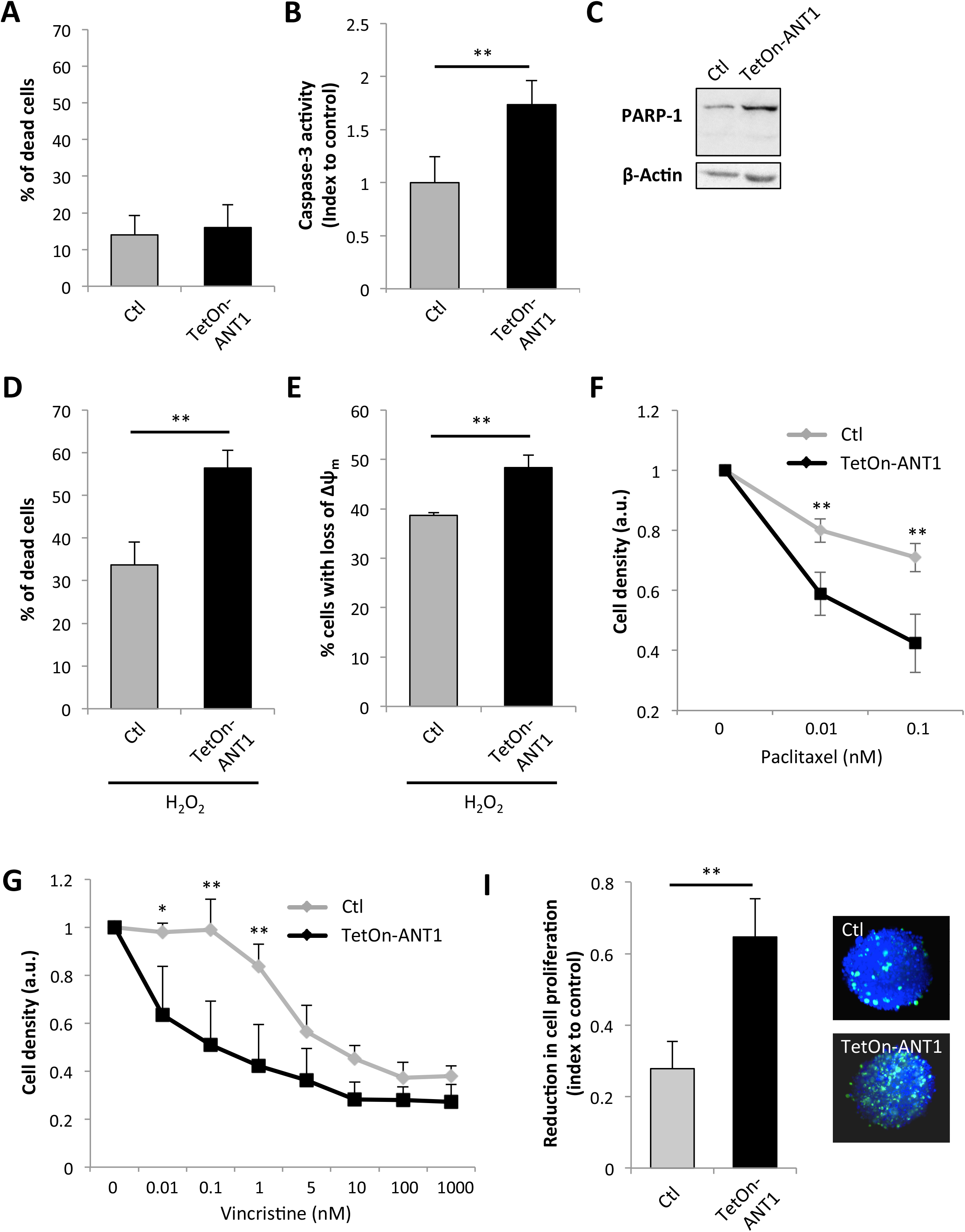
Restoration of ANT1 is sufficient to sensitize RMS cells to death induced by stress and chemotherapies. **a.** Absence of impact of ANT1 expression restoration using an inducible Tet-On system (ANT1^High^ ACC-489 cells) on percentage of dead cells. Dead cells were stained by DAPI and quantified using Nuclecounter system. Results are presented as means+/−std; n=3. **b.** Increase in basal level of caspase-3 activity in ANT1^High^ ACC-489 cells. Results are presented as means+/−std; n=3. **: p<0.01; two-sided independent samples T-test. **c.** Absence of PARP cleavage in ANT1^High^ ACC-489 cells in normal growth conditions. Actin is used as a loading control. **d-e.** Increase in percentage of dead cells (d) and with loss of Δψ_m_ (e) in ANT1^High^ ACC-489 cells upon H_2_O_2_ treatment, compared to control cells. Quantification was made by quantification of both cell populations using respectively DAPI and JC-1 stainings (Nucleocounter). Results are presented as means+/−std; n=3. **: p<0.01; two-sided independent samples T-test. **f-g.** Increased sensitivity of ANT1^High^ ACC-489 cells to 24 hours of Paclitaxel (f) or Vincristine (g) treatment, compared to control cells (WST1 assay). Results are presented as means+/−std; n=3. *: p<0.05, **: p<0.01; two-sided independent samples T-test. **i.** Increased sensitivity of ANT1^High^ CCL-136 3D spheroids to Paclitaxel treatment, compared to control 3D structures. Left panel: CellTiter-Glo® assay. Results are presented as means+/−std; n=5. **: p<0.01; two-sided independent samples T-test. Right Panel: caspase-3 activity measured my immunofluorescence. Spheroids were imaged by SPIM after active Caspase 3 (green) and DAPI (blue) immunostainings. Representative images are shown.

Altogether, these results suggest that ANT1 could somehow prime cell to death by modulating the basal level of caspase-3 activation, and thereby impact their ability to resist to stress. Consistently, loss of ANT1 is associated with an increased resistance to stress and to conventional chemotherapies that are used for RMS treatments.

## DISCUSSION

Mitochondria are a global hub of intracellular signalisation, regulating energy production, metabolism, stress or apoptosis response to stimuli for example. ANT1 was already demonstrated to play a pivotal role in some of these processes, being involved both in control of OXPHOS metabolism and release of pro-apoptotic factors from mitochondria stroma. Here, we show that loss of ANT1 expression is sufficient to perturb significantly mitochondria functioning at different levels, thereby supporting oncogenic properties of RMS tumoral cells.

Myogenesis relies on a global reprogramming of myogenic precursors, which switch from a dormant to a proliferative state before becoming quiescent again, with acquisition of a metabolism able to sustain the high energetic cost of muscle contraction (30, 39). Moreover, it has been shown that an increased resistance to apoptosis accompanies this differentiation process.

During myogenesis, it was established that ANT1 expression is enhanced to sustain the switch from a glycolytic metabolism observed in myoblasts to an oxidative one in myofibers. We observed here that ANT1 expression is low in RMS tumors, and is notably reduced compared to fetal muscles, which are frequently used as their non-tumoral equivalents (40, 41). Since the precise origin of RMS cells is not clearly defined and may result from the oncogenic transformation of different myogenic precursors, it is not clear whether the low levels of ANT1 expression correspond to maintenance of the signature observed in early myogenic precursors or to a secondary loss in the framework of tumor transformation.

In any cases, our results support the view that increased expression of ANT1 during myogenesis is not only a consequence of muscle cells differentiation but might also impact on this process, notably by modulating caspase-3 steady-state activity, which has been shown to be a key event during myogenesis (35, 42–44). It has been shown that perturbing specific metabolic pathways impact on myogenic cell fate. We provide here some evidences that ANT1 low expression level is sufficient to drive RMS cells in an immature proliferative state, corresponding somehow to early steps of regeneration process.

We observed that low expression level of ANT1 confers selective advantages to RMS tumor cells. Few studies have been dedicated so far to the precise definition of ANT1 role in cancers, due to the high level of homology between ANT1 and ANT2, which preclude their distinction. However, apparently contradictory results have been reported so far regarding ANT1 role in tumorigenesis by the two studies published so far. Indeed, these two independent publications indicate that ANT1 silencing in glioblastoma cells (38) or its overexpression in breast cancer cells (36) both leads to reduced survival. Our results somehow reconcile these contradictory observations by showing that ANT1 indeed acts as a tumor suppressor gene, directly by priming cell to death, thereby increasing their sensitivity to stress conditions, but also indirectly by engaging cells into a differentiation process, via modulation of metabolism and modulation of death cascades notably. However, total loss of ANT1 likely has deleterious effect notably by preventing metabolic plasticity of tumor cells and by increasing the oxidative stress level upon a toxic threshold.

Thus, it likely exists an optimal low level of ANT1 expression that strengthens the oncogenic properties of tumor cells. Perturbing this tight equilibrium could appear as an appealing targeted therapeutic strategy in RMS treatment.

---

**Supplementary Figure 1: Characterization of ANT1 low expression level features in RMS**

**a.** Prognostic value of ANT1 expression level in fusion-negative RMS (n=58), corresponding to ERMS and ARMS that do not present Pax3/7-FOXO1 translocation. Cut-off between ANT1 high and low group corresponds to the 1^st^ quartile of expression. Data were obtained from E-TABM-1202. **b-c.** ANT1 expression in RMS biopsies (n=147) is correlated to two markers of proliferation (b) KI67, and (c) CDC7. R2 cancer analysis, GSE16476 dataset. **d.** Specificity of ANT1 silencing by CRISPR-Cas9 in CCL-136 cells, 48 hours after doxycycline treatment. No impact was observed on ANT2 (*SLC25A5*) expression quantified by Q-RT-PCR, relatively to *HPRT* housekeeping. Results are presented as means+/−std; n=3. **e.** Increase in viable cells concentration in ANT1 CRISPR-Cas9 CCL-136 cells (ANT1^Low^ CCL-136), 48 hours after silencing of ANT1 expression by doxycycline treatment. Results are presented as means+/−std; n=3. *: p<0.05; two-sided independent samples T-test. **f.** Efficiency of siRNA targeting ANT1 in myoblasts. Results are presented as means+/−std of 3 Q-RT-PCR, using *HPRT* as a housekeeping gene; n=3. **g.** Increase in viable myoblasts concentration, 48 hours after silencing of ANT1 using a siRNA strategy. Results are presented as means+/−std; n=3. **: p<0.01; two-sided independent samples T-test.

**Supplementary Figure 2: Impact of low level of ANT1 on proliferation and differentiation**

**a-b.** ANT1 expression in RMS biopsies (n=147) is correlated to two markers of skeletal muscle (b) Troponin T3, and (c) MYBPC1. R2 cancer analysis, GSE16476 dataset.

**Supplementary Figure 3: Efficiency of an inducible Tet-On strategy to induce ANT1 expression in RMS cells and impact on cell growth**

**a-c.** Efficiency of ANT1 induction by Tet-On system in ACC-489 cells, 48 hours after doxycycline treatment, as shown by quantification of ANT1 (*SLC25A4*) expression by Q-RT-PCR, relatively to *HPRT* housekeeping (a), immunoblot with actin as a control (b), and immunofluorescence showing colocalization with the cytochrome c mitochondrial marker (c). **d-f.** Efficiency of ANT1 induction by Tet-On system in CCL-136 cells, 48 hours after doxycycline treatment, as shown by quantification of ANT1 (*SLC25A4*) expression by Q-RT-PCR, relatively to *HPRT* housekeeping (d), immunoblot with actin as a control (e), and immunofluorescence showing colocalization with the cytochrome c mitochondrial marker (f). **g.** Increased sensitivity to death of Tet-On-ANT1 CCL-136 cells treated during 6 hours by H_2_O_2_. Results are presented as means+/−std; n=3. DAPI staining quantified by Nucleocounter. **: p<0.01; independent samples T-test. **h.** Decreased viability of ANT1 Tet-On CCL-136 cells treated during 24 hours by Paclitaxel, 24 hours after induction of ANT1 expression by doxycycline treatment, compared to control cells. Results are presented as means+/−std; n=3. *: p≤0.05; two-sided independent samples T-test.

## MATERIAL AND METHODS

### Patients and RNA samples

RMS (n=67) on one side and adult (n=5), fetal (n=9), DMD (n=5), BD (n=5) and denervated (n=5) muscle biopsies were respectively collected by Centre Léon Bérard and Hospices Civils de Lyon (France) Biological Resources Centres. Tissues banking and researches conducted were performed according to national ethical guidelines, after obtention of patients’ consents. For mRNA extraction, 10μm-thick FFPE sections were deparaffinized, lysed in 200*μ*L of ATL buffer (Qiagen) with 2*μ*L Proteinase K (Roche). 1mL of Trizol (Life Technologies) was then added to isolate total RNA, before classical purification using 200*μ*L chloroform and 500*μ*L isopropanol with 1*μ*L of Glycoblue (ThermoFisher). After washing in ethanol 75%, digestion of contaminant genomic DNA was performed 1h at 37°C by resuspending the nucleic acid pellets in 15 *μ*L of 1x buffer containing 0.25 *μ*L of 100 mM DTT, 2 *μ*L of 1M DNase and 1*μ*L of RNAsine. A new step of precipitation by isopropanol/ethanol was then performed again to eliminate DNase and RNA samples were frozen (−80°C) before use.

### 2D and 3D cell cultures and design of cell models

*In vitro* studies were conducted using the human CCL-136 ERMS cells (ATCC), and the ACC-489 ARMS cells (DSMZ). CCL-136 and ACC-489 cells were routinely maintained under standard conditions (37°C and 5% CO2 in humidified incubator) respectively in Dulbecco’s Modified Eagle’s Medium (DMEM) supplemented with 10% fetal bovine serum (FBS) or in Roswell Park Memorial Institute medium (RPMI) supplemented with 10% FBS. 3D spheroids were obtained by seeding 2.5×10^3^ ACC-489 cells with 0.5% Matrigel into 96 wells low-attachment plates for 3 days.

For stress induction, cells were exposed to 0 to 1200μm (CCL-136), 0 to 75*μ*M (ACC-489) of H_2_O_2_ for 6 hours or to Paclitaxel or Vincristine as chemotherapies for 48 hours (see concentration range in Figures). z-VAD (V116-2mg, SIGMA) was used at a concentration of 20*μ*M.

To generate stable knock-down cell lines, a conditional CRISPR-Cas9-KRAB system was used. In this system, Cas9 and KRAB from Streptococcus pyogenes are placed under the control of a tetracycline-inducible promoter in pHAGE TRE dCas9-KRAB plasmid, conferring a G418 resistance to cells. Human *SLC25A4* coding sequence (NM_001151.4, ENST00000281456.11) was cloned at BfuA1 sites in the pLKo.1-puro U6 sgRNA BfuA1 stuffer, conferring puromycine resistance to cells. Both vectors are available from Addgene. CCL-136 and ACC-489 cell lines were then transduced with both plasmids thanks to lentivirus and using lipofectamine, according to manufacturer’s instructions. Clones were then selected for stable integration during 10 days in G418 (CCL-136: 1mg/mL; ACC-489: 600*μ*g/mL) and puromycin-containing medium (CCL-136: 0.6*μ*g/mL; ACC-489: 0.4*μ*g/mL). All stable knock-down cells were further tested to check ANT1 (*SLC25A4*) expression in response to doxycycline. All the experiments presented in this manuscript were performed 48 hours after doxycycline induction.

To generate inducible stable cell lines, human *SLC25A4* coding sequence was cloned at Sal1 and Cla1 sites in the pITR plasmid (gift from Prof. Rolf Marschalek, Goethe-University of Frankfurt, Germany), a sleeping beauty-based vector allowing the doxycycline inducible expression of *SLC25A4*, using Zeocin resistance gene and red fluorescent protein (mCherry) as selection markers. CCL-136 and ACC-489 cell lines were transfected with SLC25A4-pITR or an empty plasmid as a control, and a sleeping beauty transposase expression vector (SB100X). Clones were then selected for stable integration during 10 days in zeocin-containing medium. Control and ANT1-positive clones that stably integrated those constructs were selected by flow cytometry using mCherry as a marker. All stable cells were further tested to check ANT1 (*SLC25A4*) expression in response to doxycycline. All the experiments presented in this manuscript were then performed 24 hours after doxycycline induction.

### Quantitative RT-PCR

Total RNA from cell lines was extracted using the Nucleospin RNAII kit (Macherey-Nagel) and 1*μ*g was reverse-transcribed using the iScript cDNA Synthesis kit (BioRad), according to manufacturer’s instructions. Expression of ANT1 [forward primer (5’CAAGGGGAT GCTGCCTGACC3’) and reverse primer (5’GGACTGCATCATCATTCTACG3’)] and of ANT2 [forward primer (5’CACTGCAAAGGGAATGCTTCCGG3’) and reverse primer (5’GTACATGATGTCAGTTCCTTTGCG3’)] Myogenin [forward primer (5’ TGCCATCC AGTACATCGAGC 3’) and reverse primer 5’ GCAGATGATCCCCTGGGTTG 3’)] were assessed by real-time quantitative RT-PCR on a LightCycler® 480 apparatus (Roche) using the LightCycler® 480 SYBR Green I Master Mix, according to manufacturer’s instructions (Roche). The ubiquitously expressed HPRT1 [forward primer (5’TGACACTGGCAAAACAATGCA3’) and reverse primer (5’GGTCCTTTTCACCAGCAAGCT3’)] was used as an internal calibrator.

### Measurements of cell viability and density

Cells were seeded onto 6-well-plates at a density of 1*10^5^ cells/well and treated with doxycycline to modulate ANT1 expression. NucleoCounter (Chemometec, Allerød, Denmark) was used to determine cell number and viability according to the procedure provided by the manufacturer, using a co-staining of Acridine Orange (to quantify total number of cells) and DAPI (to stain dead cells).

For WST1 assay, cells were seeded into 96-well plates at 8000 cells/well. 24h, 48 and 72h after doxycycline induction, WST1 assays were performed using Cell Proliferation Reagent WST-1 (Cat No.11644807001; Roche), according to the manufacturer’s instructions. Cells were analyzed using a spectrofluorometer (Tecan Infinite® M1000 PRO).

### Cell death assays

For mitochondrial potential measurements, cells were seeded onto 6-well-plates at a density of 1×10^5^ cells/well. 24h later, cells were treated with doxycycline, and eventually with H_2_O_2_ or chemotherapies after induction of CRISPR/Tet-On systems. Cells were then collected, washed with 1ml PBS 1X, and each sample was resuspended in 12.5μl of Solution 7 (Composition Dimethyl Sulfoxide (DMSO), JC1, Chemometech). After incubation at 37°C for 10 minutes, stained cells were centrifuged at 400g for 5 min at room temperature and the supernatant was then removed completely without disturbing the cell pellet. Cell pellet was rinsed twice in 1 mL PBS 1X and then resuspended in 0.25mL of Solution 8 (Composition DAPI dilactate, Sodium azide, Chemometech). Each population of cells stained with these different dyes was then quantified NucleoCounter NC-3000 (Chemometec).

Caspase-3 activity assay was performed using Caspase 3 Fluorimetric Assay Kit (Biovision K105-400), according to manufacturer’s instructions. In brief, 2×10^5^ cells were seeded in 6 well-plates and treated with doxycycline for 24h (inducible stable cell lines) or 48h (stable knock-down cell lines). Cells were then collected and resuspended in 55 *μ*L of cell lysis buffer and incubated on ice for 30 minutes. After centrifugation, 50*μ*L of sample was loaded into a 96-well plate and mixed with 50*μ*L of reaction buffer. Fluorescence generated by related caspase activity was quantified by fluorescent detection of free AFC after cleaved from the peptide substrate DEVD-AFC using a microplate reader (Tecan Infinite® M1000 PRO). A Bradford protein assay was performed to normalize caspase activity on the total protein amount of the sample. For 3D cultures, active caspase-3 was defined by immunostaining using Cleaved Caspase 3 (9661S, cell signalling) specific antibody. In brief, spheroids were collected, fixed and permeabilized (4% paraformaldehyde, PBS1x/Triton 0.1%) blocked in PBS BSA 3% for 45 minutes. Primary antibody was added (diluted 1/200 in PBS1x) over night at +4°C. Secondary antibody was added (diluted 1/500 in PBS 1x) for an hour with DAPI at RT.

### Western-blot & Co-immunofluorescence

Cells were lysed in 500*μ*L of a pH7.6 solution containing 50mM Hepes-Sodium, 150mM NaCl, 5mM EDTA, 0.1% NP40 in the presence of protease inhibitors (Roche). Protein extracts were then analysed by immunoblot. Briefly, proteins were loaded onto 10% SDS-polyacrylamide gels and blotted onto PVDF sheets (BioRad) using TurboBlot technology (BioRad). Filters were blocked with 10% non-fat dried milk and 5% BSA in PBS/0.1% Tween 20 (PBS-T) for one hour and then incubated overnight with anti-ANT1 (sc-9299, Santa Cruz), anti-ERK1/2 (9102, Cell signaling), anti-mTOR (2983, Cell signaling), anti-FOXO3 (2497, Cell signaling), anti-Caspase9 (9502S, Cell signaling), anti-PARP (9542, Cell signaling), anti-Tubulin (ab15246, Abcam) or anti-β-actin (mAB1501R, Sigma). After three washes with PBS-Tween 0.1%, filters were incubated with the appropriate HRP◻conjugated secondary antibody (1:5000, Jackson ImmunoResearch) for 1 h. Detection was performed using ECL Chemiluminescence System (Pierce). Membranes were imaged on the ChemiDoc Touch Imaging System (Biorad).

Expression of ANT1 and cytochrome c was assessed by co-immunofluorescence on cells fixed 20 minutes in 4% paraformaldehyde and permeabilized in PBS1x/Triton0.2%, using specific antibodies (respectively sc-9299, Santa Cruz; 1:500 and ab90529, Abcam, 1:500). Fluorescence labelling were obtained using corresponding secondary antibodies coupled to Alexa Fluor-488 or Alexa Fluor-647 at a dilution of 1:500 (Invitrogen).

### Metabolic analyses

Metabolomic analyses were carried by HRMAS-NMR (High Resolution Magic Angle Spinning-Nuclear Magnetic Resonance) spectroscopy, after washing of 1×10^6^cells with D2O (Deuterium Oxide, Cortecnet) and inclusion in dedicated inserts (Cortecnet), as described previously.

Simultaneous multiparameter metabolic analysis of live cells was performed in the Seahorse XF24^®^extracellular flux analyzer (Seahorse Bioscience, USA). ACC-489 and CCL-136 cells were seeded in XF24 V7 multi-well plates (15,000 cells per well) for 5 h at 37 °C in 5% CO_2_. One hour before recording the glycolytic activity, cell culture medium was replaced with minimal DMEM (0 mM glucose) without phenol red supplemented with 143 mM NaCl, 2 mM glutamine and 1 mM sodium pyruvate, pH 7.4. Extracellular acidification rate (ECAR) was measured under these basal conditions and after sequential injections of glucose (10 mM), of the ATP synthase inhibitor oligomycin (1 μM), and of the glycolysis inhibitor 2-deoxyglucose (100 mM). To record the mitochondrial activity, the same assay medium was used and supplemented with 1 mM sodium pyruvate and 10 mM glucose. Oxygen consumption rate (OCR) was analyzed before and after sequential injections of oligomycin (1 μM), of the electron transport chain uncoupler FCCP (1 μM) and of specific inhibitors of the mitochondrial respiratory chain antimycin A/rotenone (0.5 μM). To normalize OCR and ECAR data to cell number, cells were fixed with glutaraldehyde 1%, stained with violet crystal 0.1% in methanol 20% (Sigma-Aldrich), which was finally solubilized in DMSO to measure dye absorption with a microplate spectrophotometer (600 nm). Spare respiratory capacity is calculated as the difference between maximal respiration and basal respiration as following: ((OCR after FCCP injection)−(OCR after oligomycin injection))×100. Basal extracellular acidification is calculated as the difference between glycolysis and non-glycolytic acidification as following: ((ECAR after glucose injection)−(ECAR prior glucose injection))×100.

### Detection of ROS using Cell Rox staining

CellROX® green oxidative stress reagent (Invitrogen, C10444), which is non-fluorescent while in a reduced state and exhibits a strong fluorogenic signal upon oxidation was used to detect Reactive Oxygen Species (ROS). 5×10^3^ cells were seeded in 96-well plates and treated with 1000μM H_2_O_2_, after 48 hours doxycycline induction. CellROX® green was then added in each well at a final concentration of 500ng/mL and incubated for 15min. Fluorescence intensity was then monitored with an Incucyte ZOOM® system (Essen BioScience, Michigan, USA). Phase images were obtained every two hours for 24h.

## BIBLIOGRAPHY

1. Marshall GM, Carter DR, Cheung BB, Liu T, Mateos MK, Meyerowitz JG, et al. The prenatal origins of cancer. Nature reviews Cancer. 2014;14(4):277–89.

2. Filbin M, Monje M. Developmental origins and emerging therapeutic opportunities for childhood cancer. Nature medicine. 2019;25(3):367–76.

3. Schwartz LM, Gao Z, Brown C, Parelkar SS, Glenn H. Cell death in myoblasts and muscles. Methods Mol Biol. 2009;559:313–32.

4. Brown JM, Attardi LD. The role of apoptosis in cancer development and treatment response. Nature reviews Cancer. 2005;5(3):231–7.

5. Fulda S. Cell death pathways as therapeutic targets in rhabdomyosarcoma. Sarcoma. 2012;2012:326210.

6. Saab R, Spunt SL, Skapek SX. Myogenesis and rhabdomyosarcoma the Jekyll and Hyde of skeletal muscle. Curr Top Dev Biol. 2011;94:197–234.

7. Heinicke U, Haydn T, Kehr S, Vogler M, Fulda S. BCL-2 selective inhibitor ABT-199 primes rhabdomyosarcoma cells to histone deacetylase inhibitor-induced apoptosis. Oncogene. 2018;37(39):5325–39.

8. Chamberlain JS, Metzger J, Reyes M, Townsend D, Faulkner JA. Dystrophin-deficient mdx mice display a reduced life span and are susceptible to spontaneous rhabdomyosarcoma. FASEB J. 2007;21(9):2195–204.

9. Fanzani A, Monti E, Donato R, Sorci G. Muscular dystrophies share pathogenetic mechanisms with muscle sarcomas. Trends Mol Med. 2013;19(9):546–54.

10. Fernandez K, Serinagaoglu Y, Hammond S, Martin LT, Martin PT. Mice lacking dystrophin or alpha sarcoglycan spontaneously develop embryonal rhabdomyosarcoma with cancer-associated p53 mutations and alternatively spliced or mutant Mdm2 transcripts. Am J Pathol. 2010;176(1):416–34.

11. Bauer MK, Schubert A, Rocks O, Grimm S. Adenine nucleotide translocase-1, a component of the permeability transition pore, can dominantly induce apoptosis. The Journal of cell biology. 1999;147(7):1493–502.

12. Halestrap AP. Mitochondrial permeability: dual role for the ADP/ATP translocator? Nature. 2004;430(7003):1 p following 983.

13. Kaukonen J, Juselius JK, Tiranti V, Kyttala A, Zeviani M, Comi GP, et al. Role of adenine nucleotide translocator 1 in mtDNA maintenance. Science. 2000;289(5480):782–5.

14. Stepien G, Torroni A, Chung AB, Hodge JA, Wallace DC. Differential expression of adenine nucleotide translocator isoforms in mammalian tissues and during muscle cell differentiation. The Journal of biological chemistry. 1992;267(21):14592–7.

15. Brenner C, Subramaniam K, Pertuiset C, Pervaiz S. Adenine nucleotide translocase family: four isoforms for apoptosis modulation in cancer. Oncogene. 2011;30(8):883–95.

16. Halestrap AP, Brenner C. The adenine nucleotide translocase: a central component of the mitochondrial permeability transition pore and key player in cell death. Current medicinal chemistry. 2003;10(16):1507–25.

17. Vyssokikh MY, Brdiczka D. The function of complexes between the outer mitochondrial membrane pore (VDAC) and the adenine nucleotide translocase in regulation of energy metabolism and apoptosis. Acta biochimica Polonica. 2003;50(2):389–404.

18. King MS, Thompson K, Hopton S, He L, Kunji ERS, Taylor RW, et al. Expanding the phenotype of de novo SLC25A4-linked mitochondrial disease to include mild myopathy. Neurology Genetics. 2018;4(4):e256.

19. Korver-Keularts IM, de Visser M, Bakker HD, Wanders RJ, Vansenne F, Scholte HR, et al. Two Novel Mutations in the SLC25A4 Gene in a Patient with Mitochondrial Myopathy. JIMD reports. 2015;22:39–45.

20. Tosserams A, Papadopoulos C, Jardel C, Lemiere I, Romero NB, De Lonlay P, et al. Two new cases of mitochondrial myopathy with exercise intolerance, hyperlactatemia and cardiomyopathy, caused by recessive SLC25A4 mutations. Mitochondrion. 2018;39:26–9.

21. Echaniz-Laguna A, Chassagne M, Ceresuela J, Rouvet I, Padet S, Acquaviva C, et al. Complete loss of expression of the ANT1 gene causing cardiomyopathy and myopathy. Journal of medical genetics. 2012;49(2):146–50.

22. Chevrollier A, Loiseau D, Reynier P, Stepien G. Adenine nucleotide translocase 2 is a key mitochondrial protein in cancer metabolism. Biochimica et biophysica acta. 2011;1807(6):562–7.

23. Sharaf el dein O, Mayola E, Chopineau J, Brenner C. The adenine nucleotide translocase 2, a mitochondrial target for anticancer biotherapy. Current drug targets. 2011;12(6):894–901.

24. Shi N, Xie WB, Chen SY. Cell division cycle 7 is a novel regulator of transforming growth factor-beta-induced smooth muscle cell differentiation. The Journal of biological chemistry. 2012;287(9):6860–7.

25. Winter J, Klumpe I, Heger J, Rauch U, Schultheiss HP, Landmesser U, et al. Adenine nucleotide translocase 1 overexpression protects cardiomyocytes against hypoxia via increased ERK1/2 and AKT activation. Cellular signalling. 2016;28(1):152–9.

26. Lieu EL, Nguyen T, Rhyne S, Kim J. Amino acids in cancer. Experimental & molecular medicine. 2020;52(1):15–30.

27. Jain M, Nilsson R, Sharma S, Madhusudhan N, Kitami T, Souza AL, et al. Metabolite profiling identifies a key role for glycine in rapid cancer cell proliferation. Science. 2012;336(6084):1040–4.

28. Fortini P, Ferretti C, Iorio E, Cagnin M, Garribba L, Pietraforte D, et al. The fine tuning of metabolism, autophagy and differentiation during in vitro myogenesis. Cell death & disease. 2016;7:e2168.

29. Sonkar K, Ayyappan V, Tressler CM, Adelaja O, Cai R, Cheng M, et al. Focus on the glycerophosphocholine pathway in choline phospholipid metabolism of cancer. NMR in biomedicine. 2019;32(10):e4112.

30. Pala F, Di Girolamo D, Mella S, Yennek S, Chatre L, Ricchetti M, et al. Distinct metabolic states govern skeletal muscle stem cell fates during prenatal and postnatal myogenesis. Journal of cell science. 2018;131(14).

31. Rathbone CR, Booth FW, Lees SJ. FoxO3a preferentially induces p27Kip1 expression while impairing muscle precursor cell-cycle progression. Muscle & nerve. 2008;37(1):84–9.

32. Gopinath SD, Webb AE, Brunet A, Rando TA. FOXO3 promotes quiescence in adult muscle stem cells during the process of self-renewal. Stem cell reports. 2014;2(4):414–26.

33. Baechler BL, Bloemberg D, Quadrilatero J. Mitophagy regulates mitochondrial network signaling, oxidative stress, and apoptosis during myoblast differentiation. Autophagy. 2019;15(9):1606–19.

34. McMillan EM, Quadrilatero J. Autophagy is required and protects against apoptosis during myoblast differentiation. The Biochemical journal. 2014;462(2):267–77.

35. Fernando P, Kelly JF, Balazsi K, Slack RS, Megeney LA. Caspase 3 activity is required for skeletal muscle differentiation. Proceedings of the National Academy of Sciences of the United States of America. 2002;99(17):11025–30.

36. Jang JY, Choi Y, Jeon YK, Aung KC, Kim CW. Over-expression of adenine nucleotide translocase 1 (ANT1) induces apoptosis and tumor regression in vivo. BMC cancer. 2008;8:160.

37. Zamora M, Merono C, Vinas O, Mampel T. Recruitment of NF-kappaB into mitochondria is involved in adenine nucleotide translocase 1 (ANT1)-induced apoptosis. The Journal of biological chemistry. 2004;279(37):38415–23.

38. Lena A, Rechichi M, Salvetti A, Vecchio D, Evangelista M, Rainaldi G, et al. The silencing of adenine nucleotide translocase isoform 1 induces oxidative stress and programmed cell death in ADF human glioblastoma cells. The FEBS journal. 2010;277(13):2853–67.

39. Chen F, Zhou J, Li Y, Zhao Y, Yuan J, Cao Y, et al. YY1 regulates skeletal muscle regeneration through controlling metabolic reprogramming of satellite cells. The EMBO journal. 2019;38(10).

40. Schaaf GJ, Ruijter JM, van Ruissen F, Zwijnenburg DA, Waaijer R, Valentijn LJ, et al. Full transcriptome analysis of rhabdomyosarcoma, normal, and fetal skeletal muscle: statistical comparison of multiple SAGE libraries. FASEB journal : official publication of the Federation of American Societies for Experimental Biology. 2005;19(3):404–6.

41. Tonin PN, Scrable H, Shimada H, Cavenee WK. Muscle-specific gene expression in rhabdomyosarcomas and stages of human fetal skeletal muscle development. Cancer research. 1991;51(19):5100–6.

42. Bloemberg D, Quadrilatero J. Mitochondrial pro-apoptotic indices do not precede the transient caspase activation associated with myogenesis. Biochimica et biophysica acta. 2014;1843(12):2926–36.

43. Ikeda T, Kanazawa T, Otsuka S, Ichii O, Hashimoto Y, Kon Y. Expression of caspase family and muscle- and apoptosis-specific genes during skeletal myogenesis in mouse embryo. The Journal of veterinary medical science. 2009;71(9):1161–8.

44. Liu J, Liu J, Mao J, Yuan X, Lin Z, Li Y. Caspase-3-mediated cyclic stretch-induced myoblast apoptosis via a Fas/FasL-independent signaling pathway during myogenesis. Journal of cellular biochemistry. 2009;107(4):834–44.

