## Supplementary figures and images for "Low expression of ANT1 confers oncogenic properties to rhabdomyosarcoma tumor cells via modulating metabolism and death pathways"

### Supplemental Figures

#### Supplemental Figure 1

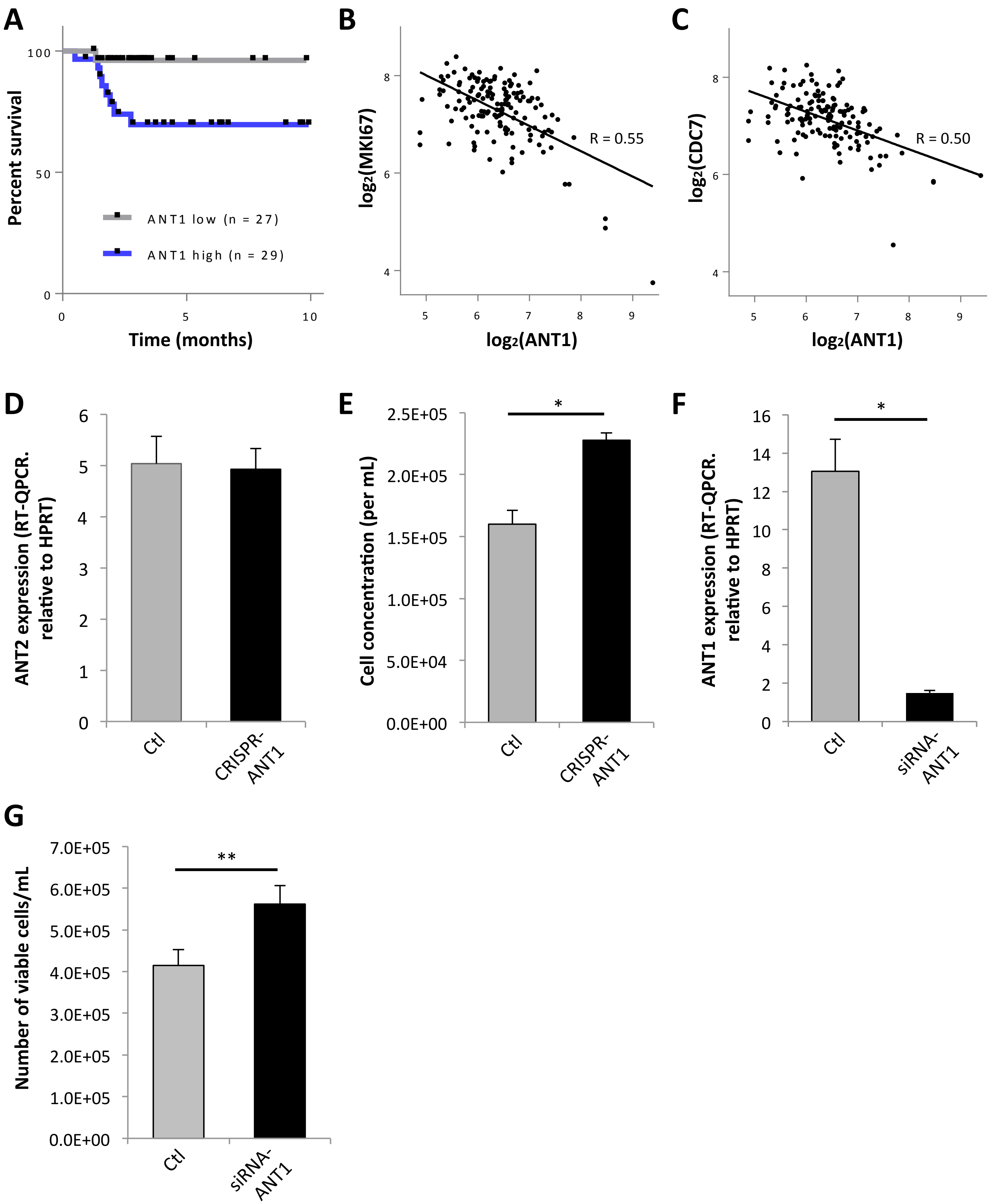

##### Supplemental Figure 2

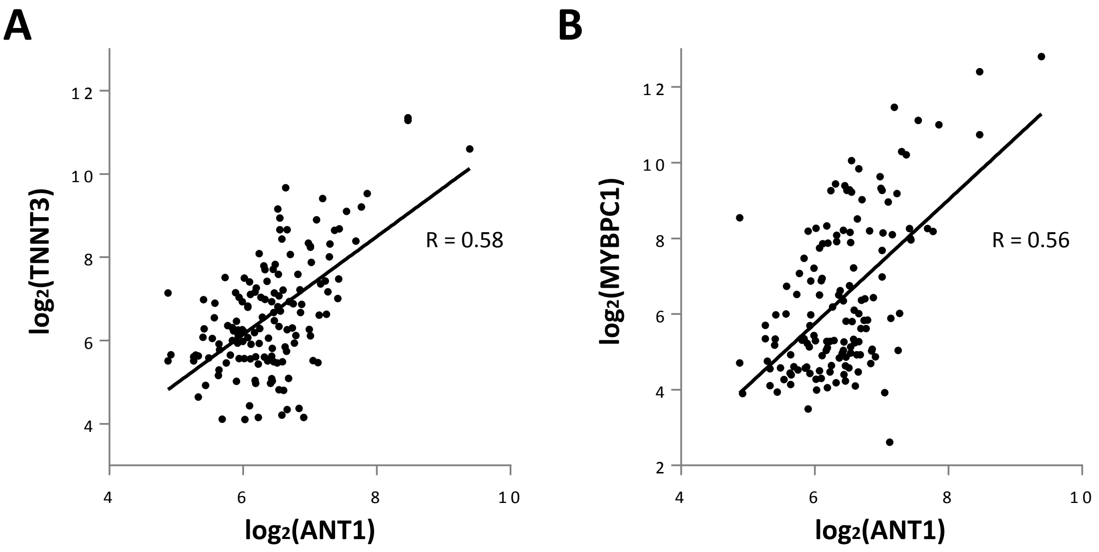

#### Supplemental Figure 3

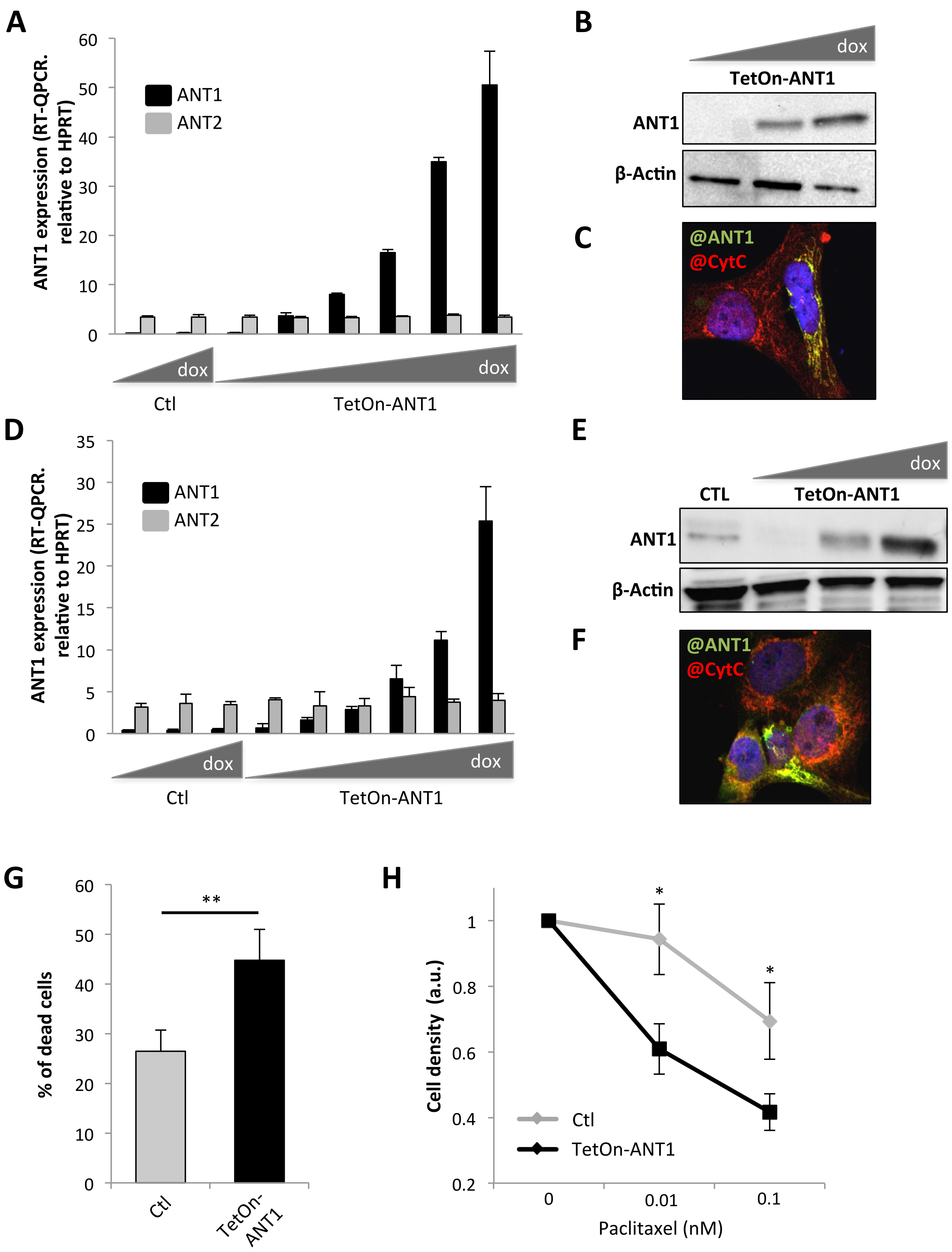
